# Promyelocytic leukemia nuclear body (PML-NB) -free intranuclear milieu facilitates development of oocytes in mice

**DOI:** 10.1101/2021.09.06.458940

**Authors:** Osamu Udagawa, Ayaka Kato-Udagawa, Seishiro Hirano

## Abstract

Promyelocytic leukemia (PML) nuclear bodies (PML-NBs), a class of membrane-less organelles in cells, are involved in multiple biological activities and are present throughout cells of adult organisms. Although the oocyte nucleus is an active region for the flux of multiple non-membranous organelles, PML-NBs have been predicted to be absent from oocytes. Here, we show that the deliberate assembly of PML-NBs during oocyte growth preferentially sequestered Small Ubiquitin-related Modifier (SUMO) protein from the nucleoplasm. SUMO not only was involved in the regulation of oocyte nuclear maturation but also was committed to the response, mediated by liquid droplet formation, to multiple stressors including nucleolar stress and proteotoxic stresses. Exogenous assembly of PML-NBs in the nucleus of oocytes affected the efficiency of the response of SUMO. These observations suggest that the PML-NB-free intranuclear milieu ensures that a reserve of SUMO remains available for emergent responses in oocyte development. This work demonstrated a benefit of the PML-NB-free intranuclear milieu, namely the ability to redirect the flux of SUMO otherwise needed to control PML-NB dynamics.

## Introduction

The nuclei of mammalian interphase cells typically contain dozens of promyelocytic leukemia (PML) nuclear bodies (PML-NBs), membrane-less organelles that are formed by phase separation. PML-NBs consist of a shell of PML protein surrounding an inner core that harbors over a hundred proteins as “client molecules”, notably including Small Ubiquitin-related Modifier (SUMO), a well-defined posttranslational modifier (Lallemand-Breitenbach and de The, 2018). The PML protein is the only resident molecule essential for the formation of PML-NBs (Lamond and Earnshaw, 1998), and polymerized PML undergoes efficient SUMOylation (Wang et al., 2018) (Li et al., 2019). Six nuclear PML isoforms have been identified in human cells; each isoform has a different C-terminal region, a distinction that results from alternative splicing of the PML-encoding primary transcript (Nisole et al., 2013). Five of the 6 isoforms (excepting PML**VI**) have a SUMO interaction motif (SIM) (Cappadocia et al., 2015), a domain through which PML can non-covalently interact with SUMO or other SUMOylated proteins that also often contain SIMs (Sahin et al., 2014). Multivalent interaction between poly-SUMO and poly-SIM is postulated as a driving force that endows PML-NBs with liquid-like properties (Banani et al., 2016). The assembly generated by mixing multivalent molecules generally shows decreased solubility, thereby promoting phase separation of the assembly; the solubility shift of resident protein is a critical factor in liquid droplet formation (Banani et al., 2017). It long has been known that specific binding of arsenic or antimony induces a sharp decline in the solubility of PML (Mu□ller et al., 1998; Hirano et al., 2015). The membrane-less properties of PML-NBs may facilitate dynamic interactions among PML-NB client molecules, which have been shown to be involved in a number of biological processes, including viral infection (Chelbi-Alix et al., 1995; Puvion-Dutilleul et al., 1995; Everett and Chelbi-Alix, 2007), DNA damage response (Louria-Hayon et al., 2003; Bernardi et al., 2004; Bøe et al., 2006), senescence (Ferbeyre et al., 2000; Pearson et al., 2000), and telomere recombination (Draskovic et al., 2009; Flynn et al., 2015).

Homozygous *Pml*^-/-^ mice have been shown to exhibit leucopenia (Wang et al., 1998) and compromised innate defense (Lunardi et al., 2011). In contrast, the phenotypes of the *Pml*^-/-^ mice during reproduction and the significance of PML in the meiosis of germ cells have not (to our knowledge) been as thoroughly studied, likely due to the normal fecundity of *Pml*^-/-^ mice (Wang et al., 1998). Indeed, PML has been demonstrated to be dispensable for embryonic development, with *Pml*^-/-^ embryos exhibiting increased resistance to acute oxidative stress compared to wild-type embryos (Niwa-Kawakita et al., 2017). These findings suggest that PML or PML-NBs play an accessory role in embryonic development. Although *Pml* mRNA is present in unfertilized mouse oocytes (Ebrahimian et al., 2010; Cho et al., 2011), the appearance of PML-NBs has not been reported, raising the possibility that *Pml* mRNA may be dormant (i.e., transcripts are stable and left untranslated). Alternatively, although PML-NBs are known to be ubiquitously distributed in adult organisms (Goddard et al., 1995; Bernardi and Pandolfi, 2003), PML protein may not be phase-separated in a manner sufficient to form PML-NBs in oocytes.

Mouse oocytes undergo growth in follicles until these cells are ready for hormone-dependent ovulation. In response to hormone signaling, fully grown germinal vesicle (GV) oocytes resume meiotic division characterized by GV breakdown (GVBD) and subsequent polar body extrusion before fertilization with sperm (Racki and Richter, 2006). During oocyte growth at meiosis prophase I, as in interphase in somatic cells, the nucleus is surrounded by a nuclear membrane and includes actively transcribing, three-compartmented nucleoli (Fulka et al., 2020). As oocyte growth proceeds, non-membranous organelles dynamically change their characteristics and their fates. For instance, processing bodies (P-bodies) disappear early in oocyte growth, with some P-body components transiently forming subcortical aggregates (SCAs; storage compartments for maternal mRNAs) in GV oocytes (Flemr et al., 2010). Decreased transcription in oocytes results in a reduced number of enlarged nuclear speckles, which appear to retain unspliced pre-mRNAs (Ihara et al., 2008). The nucleolus loses sub-compartments upon gradual shutdown of rRNA synthesis, forming nucleolus-like bodies (NLBs) surrounded by heterochromatin in the nuclei of GV oocytes (Bouniol-Baly et al., 1999). These active fluxes of non-membranous organelles spatiotemporally regulate maternal RNA metabolism during oocyte growth.

Although the oocyte nucleus is an active region for phase-separated organelles, it remains unclear why oocytes are devoid of specific membrane-less organelles such as PML-NBs and Cajal bodies (Fulka and Aoki, 2016). The present study addressed the question of why the PML-NB-free intranuclear milieu of oocytes facilitates oocyte growth.

## Experimental Procedures

### Chemicals, reagents, and antibodies

Sodium arsenite (NaAsO_2_), Triton X-100, 3-isobutyl-1-methylxanthine (IBMX), and K-modified simplex optimized medium (KSOM) were purchased from Sigma (St. Louis, MO, USA). Paraformaldehyde (PFA), dimethyl sulfoxide (DMSO), bovine serum albumin (BSA), actinomycin D (AcD), and kanamycin were purchased from WAKO (Osaka, Japan). Pregnant mare serum gonadotropin (PMSG) and human chorionic gonadotropin (hCG) were purchased from Sigma and ASKA Pharmaceutical (Tokyo, Japan). Modified human tubular fluid (mHTF) medium was purchased from Kyudo (Saga, Japan). Carbobenzoxy-L-leucyl-L leucyl-L-leucinal (MG132) was purchased from Calbiochem (San Diego, CA, USA). Epoxomicin was purchased from Enzo Life Sciences (Farmingdale, NY, USA). α-minimum essential media (MEM) medium, penicillin/streptomycin, and 4-(2-hydroxyethyl)-1-piperazineethanesulfonic acid (HEPES) were purchased from Gibco/Thermo Fisher Scientific (Grand Island, NY, USA). Lipofectamine LTX-Plus reagents, TOP10 competent cells, NuPAGE 4-12% Bis-Tris gels, lithium dodecyl sulfate (LDS) sample buffer, Bolt^TM^ WB systems, and MagicMark^TM^XP size standards were purchased from Invitrogen/Thermo Fisher Scientific (Carlsbad, CA, USA). Mineral oil was purchased from Nacalai Tesque (Kyoto, Japan). ML-792 was purchased from MedKoo Biosciences (Morrisville, NC, USA). Hoechst dye was purchased from Dojin Chemical (Kumamoto, Japan). DNase was purchased from Ambion/Thermo Fisher Scientific (Carlsbad, CA, USA). Radioimmunoprecipitation (RIPA) lysis solutions (containing 0.1% sodium dodecyl sulfate), protease inhibitor cocktail, phenylmethylsulfonyl fluoride (PMSF), and sodium orthovanadate (SOV) were purchased from Santa Cruz Biotechnology (Santa Cruz, CA, USA). KOD-FX and KOD-plus Mutagenesis Kits were purchased from TOYOBO (Osaka, Japan). Fetal bovine serum (FBS) was purchased from Biowest (Nuaille, France). ECL^TM^ Prime was purchased from GE Healthcare (Buckinghamshire, UK). The following antibodies were used in this study: anti-α thalassemia/mental retardation syndrome X-linked protein (ATRX), anti-death domain-associated protein (DAXX), anti-human PML (sc-966), Alexa 488-conjugated anti-SUMO2/3, anti-lamin B, horseradish peroxidase (HRP) -conjugated goat anti-mouse or -rabbit immunoglobulin G (IgG) (Santa Cruz Biotechnology), anti-human PML (A301-167A: Bethyl, Montgomery, TX, USA), anti-mouse PML (#05-718: Millipore), anti-SUMO1 (#4940), anti-survivin, anti-early endosome antigen 1 (EEA1) (Cell Signaling Technology, Danvers, MA, USA), anti-multi ubiquitin (Ub, #D058-3), anti-SUMO2/3 (#M114-3: MBL, Nagoya, Japan), anti-calcium-responsive transactivator (CREST) protein human antiserum (Immuno Vision, Springdale, USA), anti-synaptonemal complex protein 3 (SYCP3), anti-SUMO-conjugating enzyme (Ubc9), anti-proteasome 20*S* alpha 1+2+3+5+6+7 (20S) (Abcam, Cambridge, UK), anti-70-kDa heat shock proteins (Hsp70) (StressMarq Biosciences, Victoria, Canada), anti-splicing component, 35 kDa (SC35) (S4045: Sigma), and Alexa 488-, 594-, or 647-conjugated secondary antibodies (Molecular Probes/Thermo Fisher Scientific). The #M114-3 clone of anti-SUMO2/3 antibody also was used to generate additional Alexa 488-conjugated anti-SUMO2/3 antibody, which was labeled using an Alexa Fluor^TM^ 488 Antibody Labeling Kit (Molecular Probes/Thermo Fisher Scientific, Eugene, OR, USA)

### Collection and culture of oocytes, zygotes, and embryos

All animal procedures and protocols were in accordance with the *Guidelines for the Care and Use of Laboratory Animals* (June 2021 edition) and were approved by the Animal Care and Use Committee of the National Institute for Environmental Studies (NIES) (Approval No. 21-002). C57BL/6J mice were purchased from CLEA-Japan (Kawasaki, Japan). The animals were housed under a 12-hr/12-hr light/dark cycle with free access to food and water. Unless otherwise mentioned, oocytes were cultured in medium for *in vitro* maturation (basal medium: α-MEM medium supplemented with penicillin/streptomycin (100 units/mL and 100 μg/mL, respectively) and heat-inactivated FBS (10%)). Actively transcribed and meiotically incompetent oocytes collected from postnatal days 12-16 are referred to as “maturing oocytes” in the present study. Subsets of oocytes in the ovary form follicles that then grow (folliculogenesis), a process that initiates just before birth and continues to postnatal days 22-24 (Racki and Richter, 2006; Rodriguez et al., 2019). The status of the full-sized oocytes remains arrested at meiotic prophase with a large nucleus, a structure that is designated the germinal vesicle (GV). The GV oocyte stage can be subdivided into the non-surrounded nucleolus (NSN) stage and the surrounded nucleolus (SN) stage based on maturity. The SN stage is characterized by a transcriptionally inactive heterochromatin rim (i.e., with heterochromatin surrounding the nucleolus-like body (NLB)), wherein the cell is primed for breakdown of nuclear membrane (GVBD), a hallmark of the resumption of meiosis. Except for the oocytes depicted in Fig. S1B-D (which represent samples collected directly from adult ovaries), Metaphase I (MI) and Metaphase II (MII) -stage oocytes were obtained by *in vitro* culture as follows. For collection of fully grown GV oocytes, follicles on the ovarian surface were mechanically ruptured with a pair of forceps. GV oocytes initially were isolated in basal medium supplemented with HEPES (12.5 mM) and IBMX (0.1 mM); IBMX is a phosphodiesterase inhibitor that inhibits/controls spontaneous meiotic resumption. After IBMX was removed by washing with basal medium, oocytes were incubated for 4 hr to allow GVBD to proceed. Development of oocytes was further verified by observation of the extrusion of the first polar body (Pb1; a marker of MII-stage oocytes that have the potential to be fertilized with sperm) after an additional incubation of up to 16 hr. We regarded Pb1-free oocytes with no GV as MI-stage oocytes. To obtain embryos, female mice were initially primed by intraperitoneal injection of PMSG followed 48 hr later by injection with hCG. Superovulated oocytes were collected from the oviducts of euthanized mice by gently teasing apart of the ampulla with a 21-gauge needle (TERUMO, Tokyo, Japan) to release cumulus-oocyte complexes in mHTF medium; these oocytes then were inseminated with pre-capacitated sperm. To assess pronuclear (PN) stages after insemination, we evaluated fertilized oocytes based on the distance between the male and female pronuclei. PN stages were defined as follows: PN3, smaller pronuclei that are distributed distantly; PN5, bigger pronuclei that overlap. After fertilization, embryos were cultured further until the indicated embryonic stages or time points using KSOM medium. Cultures were performed in a drop of medium (30-80 µL) under a mineral oil overlay at 37 °C in a humidified atmosphere of 5% CO_2_ (APM-30D; ASTEC, Fukuoka, Japan), except for heat shock (HS) experiments, which were performed at 42 °C in ambient air (FMC-1000; EYELA, Tokyo, Japan).

### Western blot analysis for PML and SUMO in CHO-K1 cells

Using Lipofectamine LTX-Plus reagents, CHO-K1 cells were transiently transfected with plasmids harboring *Pml**VI*** (human PML transcript variant 5, NM_033244) or a cDNA encoding a SUMOylation-deficient version of the protein. The former construct, which we designated v5-hPML**VI**, was obtained from OriGene (Rockville, MD, USA). The latter construct, which we designated v5-hPML**VI** (K160, 490R), encodes a mutant version of the same protein in which the nucleic acid sequences encoding the K160 and K490 amino acid residues were altered to encode Arg residues (by mutation using the KOD-plus Mutagenesis Kit according to the manufacturer’s instructions). The K160 and K490 residues correspond to sites known to be required for SUMOylation (Lallemand-Breitenbach et al., 2008). The transfected cells were washed with phosphate-buffered saline (PBS, pH 7.1-7.7) and then lysed on ice with 100 µL of RIPA lysis buffer pre-mixed with protease inhibitor cocktail (1:100, vol/vol), PMSF (a serine protease inhibitor; 2 mM), and SOV (a phosphatase inhibitor; 1 mM). The lysate was centrifuged at 9,000 x g for 5 min at 4 °C and the resulting supernatant was designated as the RIPA-soluble fraction. The pellet was washed again with PBS and then treated with DNase (40 U/mL) to reduce the viscosity. Pellets were ruptured by ultrasonication at 4 °C for 30 min using a Bioruptor^TM^ (UCD-250, H-amplitude, repeating 15-sec sonication at 45-sec intervals; Cosmobio, Tokyo, Japan) followed by a 1-hr incubation at 25 °C with intermittent vortexing using a thermomixer (Eppendorf, Wesseling-Berzdorf, Germany). Aliquots were designated as the RIPA-insoluble fraction and subsequently mixed with LDS sample buffer and boiled at 95 °C for 3 min before being stocked frozen at -30 °C. Before loading onto gels, samples again were boiled at 95 °C for 10 min. Proteins in the samples were resolved by LDS-polyacrylamide gel electrophoresis (LDS-PAGE) and electroblotted to membranes; the membranes were blocked and then probed with primary antibodies overnight at 4 °C before hybridization with secondary antibodies for 1 hr at room temperature. Signals obtained with ECL^TM^ Prime Chemiluminescent HRP Substrate were detected with an FAS1100 (TOYOBO).

### Plasmid or mRNA microinjection

The Emerald Green Fluorescent Protein (EmGFP) -encoding region from Vivid Colors^TM^ cDNA pcDNA^TM^6.2/N-EmGFP-DEST Vector (Thermo Fisher Scientific) was placed upstream of the coding sequence of *Pml**VI*** (human PML transcript variant 5, NM_033244) as described previously (Hirano et al., 2018). The fusion protein encoded by the resulting construct was designated GFP-hPML**VI**. Each plasmid, including v5-hPML**VI** and v5-hPML**VI** (K160, 490R), was diluted to a concentration of 5 ng/µL with Milli-Q water. mRNA preparation was conducted by BEX (Tokyo, Japan) by *in vitro* transcription from v5-hPML**VI** or v5-hPML**VI** (K160, 490R) using a HiScribe^TM^ T7 ARCA mRNA Kit with tailing (New England Biolabs, Ipswich, MA, USA) and a MEGAclear^TM^ Transcription Clean-Up Kit (Thermo Fisher Scientific). Each mRNA was diluted to a concentration of 25 or 100 ng/µL with RNase free water (Jena Bioscience, Jena, Germany). Each solution was loaded into a DNA injection pipette (TIP-DNA [LIC-OD1], NAKA Medical, Tokyo, Japan), and the pipettes were placed in an IM-9B microinjector (Narishige, Tokyo, Japan). MN-4 and MMO-202ND manipulators (Narishige) were adapted to an IX70 inverted microscope (Olympus, Tokyo, Japan) via NO-PIX-4-P (Narishige). Plasmid or mRNA solutions were injected into each oocyte nucleus or cytoplasm (respectively) until adequate swelling of the nuclear or plasma membrane (respectively) was observed. Injection was conducted in the basal medium supplemented with HEPES (12.5 mM) and IBMX (0.1 mM). Injected oocytes were washed and cultured without HEPES during the interval indicated for each experiment.

### Immunofluorescent (IF) staining

Except for the live imaging of GV oocytes in Figure 2A, oocytes, zygotes, and embryos at each indicated stage were fixed with 4% phosphate-buffered PFA (pH 7.0-7.4) at room temperature. Cells were permeabilized with 0.5% Triton X-100 in PBS, blocked with 5% BSA-PBS, and then stained with the indicated primary antibodies in 1% BSA-PBS. Cells were visualized with Alexa-conjugated secondary antibodies; Hoechst dye was included to stain DNA. Cells were placed in small drops (4 µL each) of 1% BSA-PBS, covered with mineral oil, in a glass-bottomed 35-mm petri dish (AGC techno glass, Shizuoka, Japan). Images were captured by confocal microscopy (Leica TCS-SP5; Leica, Solms, Germany).

For the detection of nascently translated polypeptides, oocytes or embryos were cultured with O-propargyl-puromycin (OP-puro) (obtained as part of a Click-iT^®^ Plus OPP Protein Synthesis Assay Kit; Molecular Probes/Thermo Fisher Scientific) at a concentration of 20 µM for the indicated time periods, followed by Click reaction according to the manufacturer’s instructions. Images were captured by confocal microscopy.

As a reference for PML-NB staining, bone marrow from euthanized male C57BL/6J mice was exposed by removing both ends of the femur; bone marrow cells then were obtained by flushing the marrow with PBS using a 25-gauge needle (TERUMO). The resulting cells were immediately cytocentrifuged onto slide glass and air-dried. The cell preparations were used for immunostaining for visualization of endogenous PML-NBs. Images were captured by fluorescence microscopy (ECLIPSE 80i; Nikon, Tokyo, Japan).

### Electron microscopic (EM) analysis

GV oocytes were cultured for 20 hr. Metaphase oocytes were fixed in 4% PFA, 0.2% glutaraldehyde, and 0.5% tannic acid in 0.1 M cacodylate buffer, pH 7.4, for 100 min at room temperature. The fixed oocytes then were washed with 0.1 M cacodylate buffer. Further procedures were conducted by Tokai Electron Microscopy, Inc. (Nagoya, Japan). After dehydration, the sample was embedded in resin (LR White; London Resin, Berkshire, UK). The polymerized resin was ultra-thin sectioned at thicknesses of 80 nm, and the sections were mounted on nickel grids. The grids were incubated with the primary antibody (anti-mouse PML, #05-718) in 1% BSA-PBS overnight at 4 °C. The grid-mounted sections subsequently were incubated for 1 hr at room temperature with the secondary antibody conjugated to 20-nm gold particles (EMGMHL20; BBI Solutions, Crumlin, UK). Grids then were placed in 2% glutaraldehyde in 0.1 M cacodylate buffer. Subsequently, the grids were dried and stained with 2% uranyl acetate for 15 min, and then placed in lead staining solution (Sigma) at room temperature for 3 min. The grids were observed by transmission electron microscopy (JEM-1400Plus; JEOL, Tokyo, Japan).

### Data analysis

Where indicated, data are presented as means with the standard error of the mean (SEM).

## Results

### PML-NBs are not formed during development of oocytes

To understand the location of PML protein in oocytes during and after the resumption of meiosis, GV oocytes were collected from ovaries and cultured *in vitro*. In the nucleus of GV oocytes, PML accumulated around chromatin near the nuclear membrane (Fig. 1A; **Fig. S2C**). The peri-chromosomal distribution of PML protein also was observed in oocytes in the GVBD, MI, and MII stages (Fig. 1A). Immuno-EM analysis (using anti-mouse PML primary antibody and gold-conjugated secondary antibody) revealed that gold particles (indicative of PML-positive staining) were localized peri-chromosomally as amorphous protein aggregate-like structures (**Fig. S1A**). IF analysis of PML, in combination with staining for a chromosomal passenger complex marker and a kinetochore marker, revealed that PML was arranged in a pattern extending from the kinetochores with localization along the chromosome arms (**Fig. S1B,C**). In contrast to mitotic assemblies of PML proteins (MAPPs), which are tethered to early endosomes in metaphase-stage somatic cells (Dellaire et al., 2006), PML in oocytes never co-localized with early endosomes (which themselves were labeled with anti-EEA1 antibodies, as assessed by IF) during metaphase (**Fig. S1D**). To examine the spatiotemporal distribution of PML protein, oocytes at various meiotic stages were collected. Among ovarian cells (55 of which were surveyed in detail) collected from postnatal day 1 mice, cells that stained positively for SYCP3 (a marker of germ cells in meiotic prophase) exhibited nuclear staining for PML just beneath the inner nuclear membrane; however, PML was not detected as nuclear bodies (NBs) in these cells (Fig. 1B). In contrast, NBs were seen in 98.1% of the 162 control (bone marrow) cells examined **(Fig. S1F)**. Similarly, in the pronuclei of zygotes as well as fertilized 2-cell embryos, PML protein was observed just beneath the inner nuclear membrane (Fig. 1C), where the heterochromatin is known to be enriched (Bersaglieri and Santoro, 2019). When a similar analysis was performed on zygotes, no NBs were observed in the nucleoplasm among the 32 zygotes that were surveyed, (Fig. 1C). During the development of preimplantation embryos, PML-NBs were unambiguously detected in the nucleus of each blastomere, at least in the morula to early blastocyst stage (**Fig. S1E**); In post-morula-stage embryos, PML also was slightly enriched beneath the nuclear membrane, as observed in 2-cell embryos (Fig. 1C). PML proteins in NBs exhibited co-localization with SUMO and with DAXX (**Fig. S1E**), which are representative clients of the PML-NBs (Lallemand-Breitenbach and de The, 2018). Thus, it appeared that oocyte PML is associated with the heterochromatin, a localization that is coordinated with the turnover of nuclear membrane in oocytes undergoing meiosis.

**Fig. 1.**
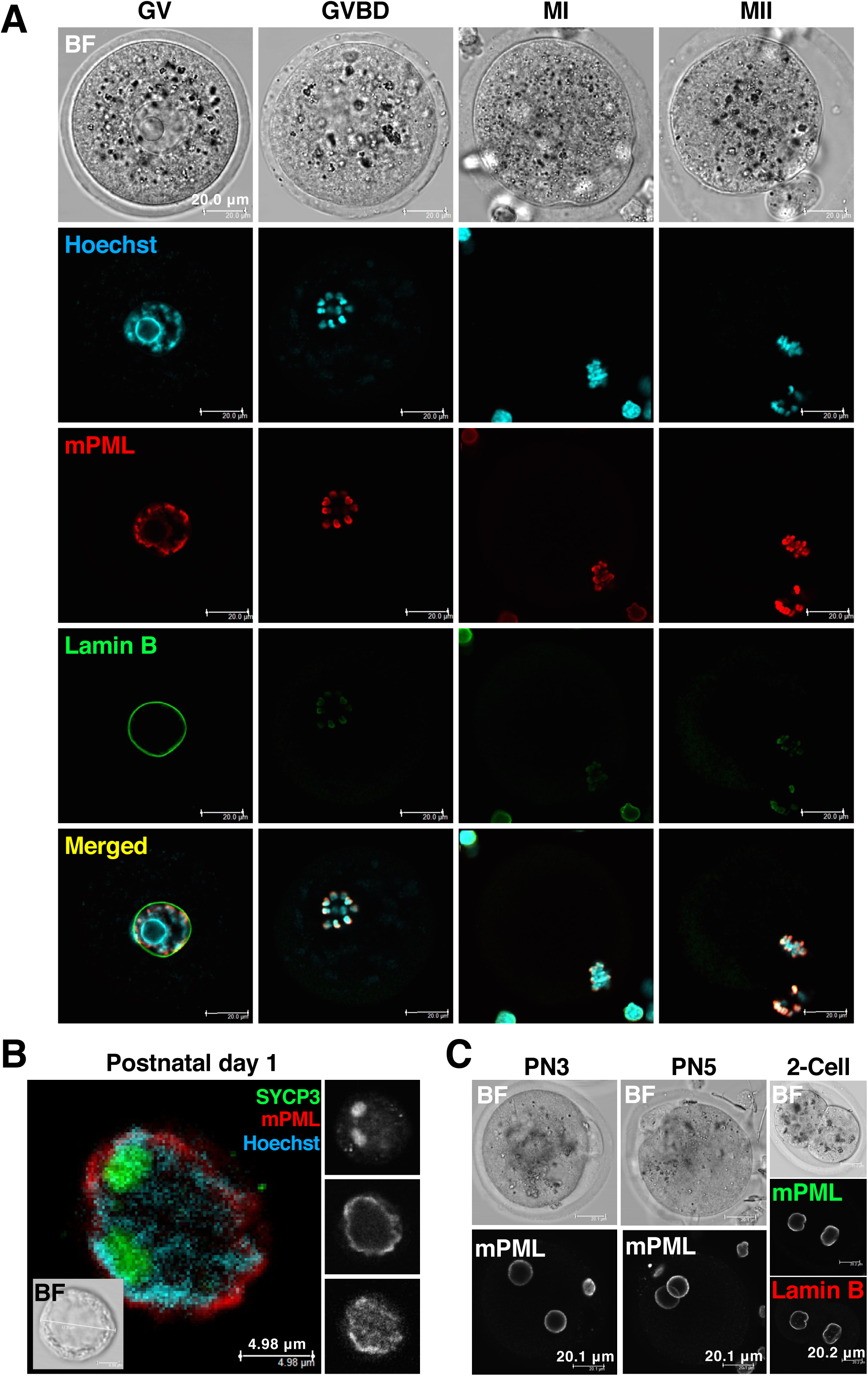
Promyelocytic leukemia (PML) nuclear bodies (PML-NBs) are not formed during oocyte development. (A) Subcellular localization of endogenous PML in oocytes during and after meiotic resumption. Germinal vesicle (GV) oocytes were cultured to obtain metaphase II (MII) -stage oocytes. Breakdown of the nuclear membrane (GV breakdown; GVBD), a hallmark of meiotic resumption, was evaluated by stereomicroscopy during culturing and verified by staining with anti-lamin B antibody (green). PML was visualized by staining with anti-mouse PML (mPML, red) antibody. BF, bright-field images of the oocytes. Scale bars, 20 μm. (B) Representative image of endogenous PML in oocytes collected from mice at postnatal day 1. The insets show separate images for each channel. Scale bars, 4.98 µm. (C) Representative images of endogenous PML in zygotes at the indicated pronuclear (PN) stages just after fertilization. The image of the pronuclear stage 5 (PN5) zygote was reconstructed as a z-stack image. The inner nuclear membrane of the 2-cell embryo also was co-labeled with anti-lamin B antibody fluorescing in a different color. BF, bright-field images of the zygotes and the embryo. Scale bars, 20.1 and 20.2 µm, respectively.

### The dynamics of nascent PML-NBs in the nucleoplasm demands SUMO

To address if any possible advantages are present in the PML NB-free intranuclear milieu of oocytes during development, we attempted the deliberate assembly of NBs in the nuclei of GV oocytes. For concise interpretation, we injected a plasmid that encodes GFP-hPML**VI**, a SUMO interaction motif (SIM) -free variant of PML, avoiding non-covalent interaction with SUMO or the SUMOylated moieties of modified proteins that often also contain SIMs. Despite a low success rate of expression (<10%), presumably due to the transcriptional quiescence of GV oocytes, injection of this construct resulted in the formation of NBs in the nuclei of GV oocytes (Fig. 2A). In these oocytes, exogenously formed GFP-hPML**VI**-induced PML-NBs did not exhibit colocalization with either the endogenous ATRX or DAXX (Fig. 2B), which are known clients of PML-NBs (Lallemand-Breitenbach and de The, 2018). In contrast, GFP-hPML**VI**-induced PML-NBs exhibited colocalization with endogenous SUMO2/3 in a manner that depended on the degree of NB formation. This dependency also was observed in actively transcribing oocytes obtained from mice at approximately postnatal day 12-16 (i.e., maturing oocytes that are meiotically incompetent (De La Fuente, 2006)) (Fig. 2C). A higher efficiency of plasmid expression (reaching >70%) was obtained in these maturing oocytes than in GV oocytes. Notably, in the presence of arsenite, even the faintly visible nascent GFP-hPML**VI**-induced PML-NBs efficiently sequestered SUMO (Fig. 2D). Given the observed tight association between GFP-hPML**VI**-induced PML-NBs and SUMO, we next assessed the effects of PML-NB formation on molecules involved in the SUMOylation-triggered PML catabolism and nuclear stress responses. Although it has been reported that ubiquitin and proteasomes are recruited to SUMOylated PML when exposed to arsenic (Lallemand-Breitenbach et al., 2008), these two molecules and a related chaperone were not recruited to GFP-hPML**VI**-induced PML-NBs (Fig. 2E). Other than the nucleolus, PML-NB also has been reported to act as overflow compartments for misfolded proteins, a process that occurs under conditions of proteotoxic stress and is mediated by the ubiquitin-proteasome system upon in the nuclei of cultured somatic cells (Uozumi et al., 2016; Mediani et al., 2019). The relative paucity of the compartments dealing with the proteinous wastes could be a disadvantage for, PML-NB-free, the oocyte. In practice, we showed that aberrant polypeptides (generated in the presence of the proteasome inhibitor MG132), which were labeled with an analog of puromycin (OP-puro), accumulated only in the nucleolus, and never in GFP-hPML**VI**-induced PML-NBs (Fig. 2F; **Fig. S2A**). Similarly, in post-morula-stage embryos under proteotoxic stress, OP-puro-labeled aberrant polypeptides did not accumulate in endogenously appearing PML-NBs (**Fig. S2B**). Together, these results suggested that oocyte nucleoplasm can accommodate PML-NBs that preferentially sequester SUMO.

**Fig. 2.**
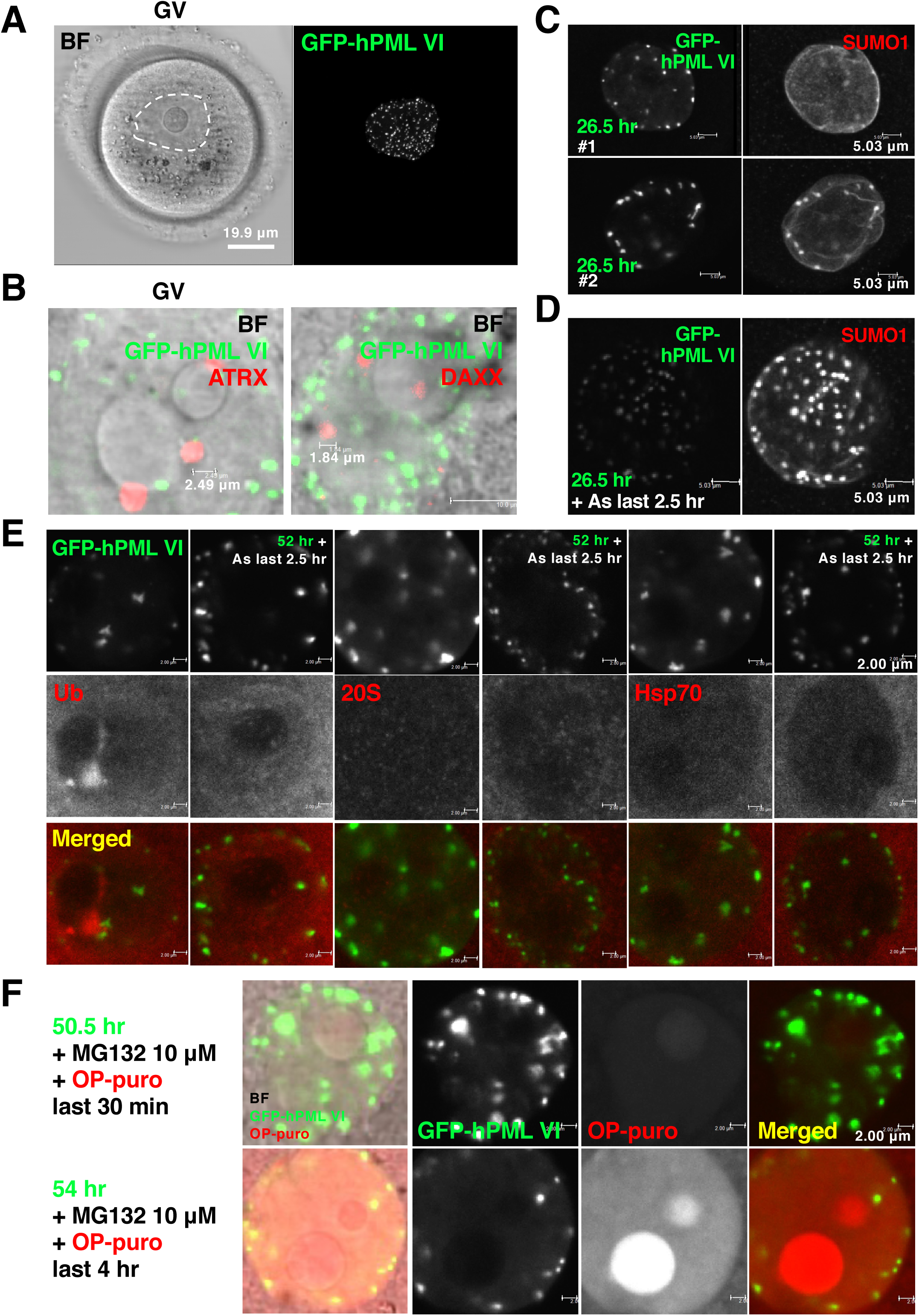
Characterization of nuclear bodies (NBs) deliberately assembled in the nuclei of oocytes. (A) Representative z-stack image of exogenously formed PML-NBs in a germinal vesicle (GV) oocyte, a model to gain insight into the benefit of a NB-free intranuclear milieu in oocytes. Briefly, GV oocytes were injected with a plasmid harboring an Emerald Green Fluorescent Protein (EmGFP) -encoding sequence upstream of the sequence of *Pml**VI*** (human PML transcript variant 5, NM_033244), and cultured for 53 hr. The resulting encoded protein was designated GFP-hPML**VI**. BF, bright-field image of the oocyte. Scale bars, 19.9 µm. (B) Representative images of exogenously formed GFP-hPML**VI**-induced PML-NBs and of representative PML-NB markers in GV oocytes. α thalassemia/mental retardation syndrome X-linked protein (ATRX) and death domain-associated protein (DAXX) are representative clients of PML-NBs. Oocytes were visualized by immunofluorescent staining with anti-human PML antibody and by co-staining with anti-ATRX (red, left) or -DAXX (red, right) antibodies. Scale bars, 2.49 and 1.84 μm, respectively. (C) Representative images of GFP-hPML**VI**-induced PML-NBs in the nuclei of actively transcribing maturing oocytes; actively transcribing maturing oocytes were used to obtain a higher efficiency of plasmid expression. Maturing oocytes were injected with a plasmid encoding GFP-hPML**VI** and cultured for 26.5 hr. Oocytes were stained with anti-human PML (green) and anti-SUMO1 (red) antibodies. (#1) While GFP-hPML**VI**-induced PML-NBs were formed, a lower degree of SUMO sequestration was observed. (#2) GFP-hPML**VI**-induced PML-NBs were fully formed and SUMO was well sequestered. Scale bars, 5.03 μm. (D) Representative z-stack image of the nucleus of a maturing oocyte injected with the plasmid encoding GFP-hPML**VI** and cultured for 24 hr followed by treatment with 3 μM arsenite (As) for 2.5 hr. Oocytes were stained with anti-human PML (green) and anti-SUMO1 (red) antibodies. Note the intensive sequestration of SUMO, even by the faintly visible GFP-hPML**VI**-induced PML-NBs. Scale bars, 5.03 μm. (E) Representative image of the nucleus of a maturing oocyte injected with the plasmid encoding GFP-hPML**VI** and cultured for 49.5 hr, then either treated with 3 μM arsenite or left untreated for another 2.5 hr. Oocytes were stained with anti-multi ubiquitin (Ub), anti-proteasome 20S alpha 1+2+3+5+6+7 (20S), and anti-70-kDa heat shock protein (Hsp70) (red) antibodies. Scale bars, 2.00 μm. (F) An examination of whether GFP-hPML**VI**-induced PML-NBs act as overflow compartments for misfolded proteins. Representative image of the nucleus of a maturing oocyte injected with the plasmid encoding GFP-hPML**VI** and cultured for 50 hr before labeling of newly synthesized aberrant polypeptides with an analog of puromycin (OP-puro, 20 μM) (red) combined with proteasome inhibitor (MG132, 10 μM) for 30 min or 4 hr. Scale bars, 2.00 μm.

We next used the deliberate assembly of PML-NBs to more thoroughly analyze SUMO availability in maturing oocytes. First, oocytes were manipulated to limit the availability of SUMO with which PML could interact. Specifically, we injected oocytes with a construct (v5-hPML**VI** (K160, 490R)) that encodes a mutated SIM-free variant of PML that has decreased SUMOylation efficiency, thereby decreasing the protein’s avidity for SUMO and the SIM moiety of SUMOylated proteins (**Fig. S3A**). We found that oocytes injected with plasmid v5-hPML**VI** (K160, 490R) exhibited a decreased number (2.22 ± 0.36 (mean ± SEM), across 12 oocytes) of enlarged v5-hPML**VI** (K160, 490R) -induced PML-NBs compared to the dozens observed in oocytes injected with the wild-type PML**VI**-encoding plasmid (Fig. 3A,B). In the v5-hPML**VI** (K160, 490R) -injected oocytes, SUMO protein accumulated with the v5-hPML**VI** (K160, 490R) -induced PML-NBs, but did not merge with the PML (K160, 490R) shell of the NB, as if the SUMO protein were physically isolated within the PML-negative inner core (Fig. 3B). While arsenite efficiently promotes SUMOylation of PML, the SUMOylation-deficient mutant of PML has been reported to remain biochemically responsive to arsenite (Lallemand-Breitenbach et al., 2001). We first confirmed this observation by showing (using our system) that arsenite induces a sharp decline in the solubility (which favors *de novo* phase separation (Banani et al., 2017)) of v5-hPML**VI** (K160, 490R) -encoded protein, similar to that seen with the wild-type v5-hPML**VI** -encoded protein (**Fig. S3B**). In oocytes injected with the plasmid v5-hPML**VI**, arsenite treatment altered the spatial relationships between PML and SUMO from co-localization (Fig. 3A) to spherical co-layering (Fig. 3C). In contrast, we observed the nascent formation of small-dot structures upon arsenite treatment in the v5-hPML**VI** (K160, 490R) -injected oocytes (Fig. 3D**#1,#2, arrows**), and found that these structures frequently were clustered (Fig. 3D**#2, asterisks**). Notably, partial scaffolding of SUMO within the inner cores of v5-hPML**VI** (K160, 490R) -induced PML-NBs assumed crooked shapes, presumably indicating partial SUMOylation of the v5-hPML**VI** (K160, 490R) -encoded protein (Fig. 3D**#1,#2, arrowheads**).

**Fig. 3.**
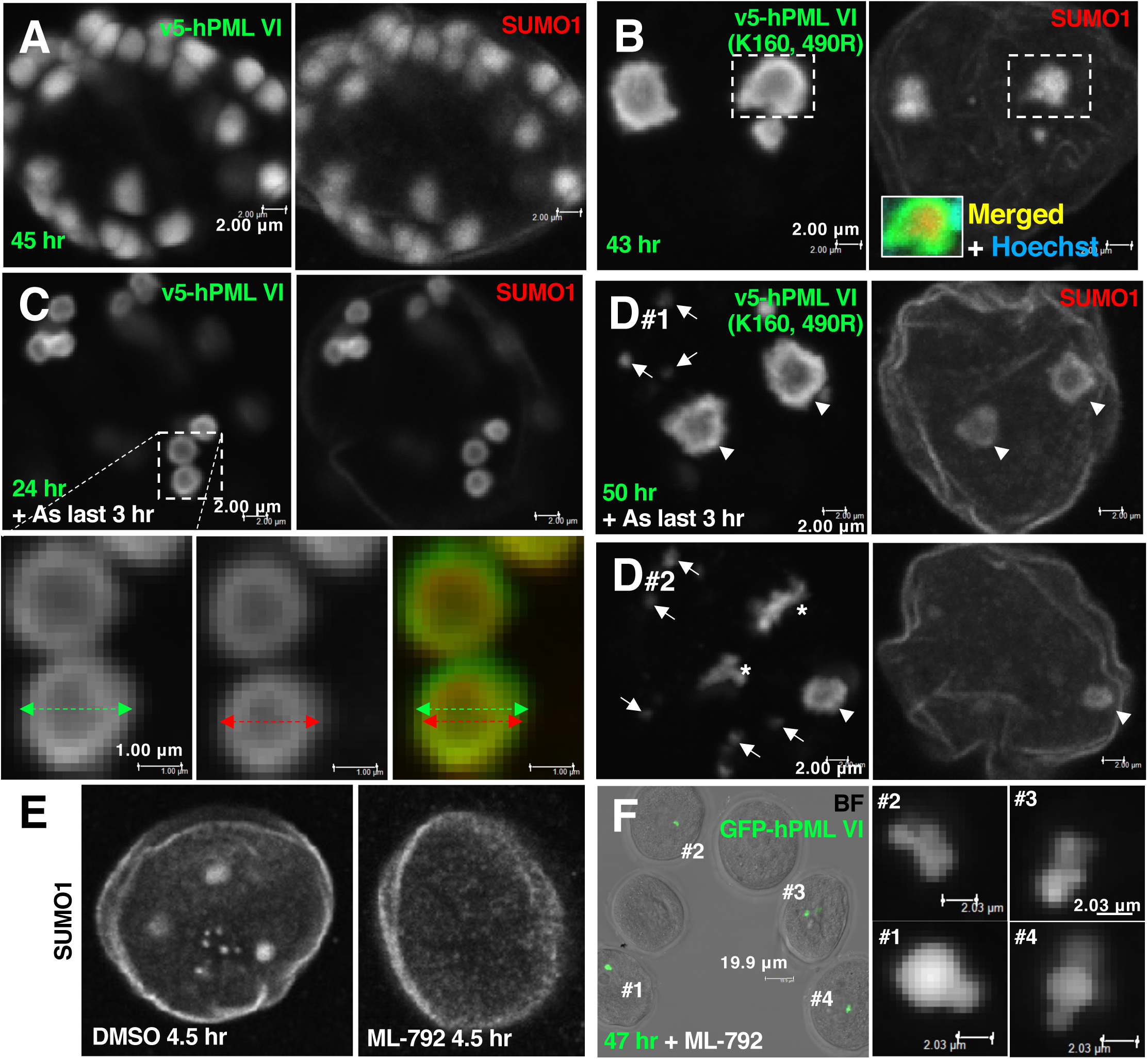
The dynamics of nascent PML-NBs in the nucleoplasm demands SUMO. (A and B) Representative images of the nuclei of maturing oocytes injected with plasmid v5-hPML**VI** (encoding wild-type human PML**VI**) (A, z-stack) or plasmid v5-hPML**VI** (K160, 490R) (encoding a human PML **VI** that has limited avidity for SUMO) (B, z-stack) and cultured for 45 hr or 43 hr, respectively. Oocytes were stained with anti-human PML (green) and anti-SUMO1 (red) antibodies. (B, inset) Merged images with Hoechst staining of (B). Scale bars, 2.00 μm. (C) Representative image of the nucleus of a maturing oocyte injected with plasmid v5-hPML**VI** and cultured for 21 hr followed by treatment with 3 μM arsenite (As) for 3 hr. Scale bars, 2.00 μm. Enlarged images of the inset with diameter information. Scale bars, 1.00 μm. (D) Representative images of the nuclei of maturing oocytes injected with plasmid v5-hPML**VI** (K160, 490R) and cultured for 47 hr (D#1 and 2, z-stack) followed by treatment with 3 μM arsenite for 3 hr. Oocytes were stained with anti-human PML (green) and anti-SUMO1 (red) antibodies. Scale bars, 2.00 μm. Arrows, nascent v5-hPML**VI** (K160, 490R)-induced PML-NBs upon arsenite treatment; arrowheads, enlarged v5-hPML**VI** (K160, 490R) -induced PML-NBs and partial scaffolding of SUMO within the inner cores of v5-hPML**VI** (K160, 490R) -induced PML-NBs; asterisks, nascent v5-hPML**VI** (K160, 490R) -induced PML-NBs that frequently are clustered upon arsenite treatment. (E) Representative z-stack images of the nuclei of maturing oocytes treated with dimethyl sulfoxide (DMSO, vehicle control) or with 20 μM ML-792, an inhibitor of a SUMO-activating enzyme limiting the nucleoplasmic availability of SUMO, for 4.5 hr. Oocytes were stained with anti-SUMO1 antibody. (F) Representative z-stack images of maturing oocytes injected with a plasmid encoding GFP-hPML**VI** and cultured in the presence of 20 μM ML-792 for 47 hr. Oocytes were stained with anti-human PML (green) antibody. Enlarged images of GFP-hPML**VI**-induced PML-NBs that clustered together are shown on the right. BF, bright-field image of the oocytes. Scale bars, 19.9 μm (insets #1 to #4, 2.03 μm).

Next, to understand the clustering of the nascent v5-hPML**VI** (K160, 490R) -induced PML-NBs, we treated maturing oocytes with ML-792 (an inhibitor of a SUMO-activating enzyme) to limit the nucleoplasmic availability of SUMO (He et al., 2017). Compared to the nuclei of maturing oocytes treated with vehicle, the nuclei of maturing oocytes treated with ML-792 lacked general staining with anti-SUMO1 antibody, exhibiting staining only at the nuclear membrane (Fig. 3E). Maturing oocytes injected with the GFP-PML-encoding plasmid and cultured in the presence of ML-792 exhibited a decreased number (1.96 ± 0.18; 27 oocytes cultured for a shorter period of 22 hr were quantified) of GFP-hPML**VI**-induced PML-NBs that clustered together (Fig. 3F). Collectively, these results suggested that maturing oocytes maintain the dynamics of PML-NBs at the cost of SUMO availability in the nucleoplasm.

### PML-NBs potentially affect the efficiency of the response of SUMO mediated by liquid droplet formation upon exposure to multiple stresses

We noticed that in maturing oocytes, a portion of the SUMO signal was detected as small drops (Fig. 4A) in the nucleolus, which is itself a droplet formed by liquid-liquid phase separation (Feric et al., 2016). Transcriptional inhibition with actinomycin D (AcD), an inducer of a nucleolar stress response (Sundqvist et al., 2009), resulted in dispersion of SUMO drops within the nucleoplasm (Fig. 4A). The nucleoplasm-localized SUMO signal following AcD exposure was sequestered by GFP-hPML**VI**-induced PML-NBs (Fig. 4B). These results suggested that formation of PML-NBs in transcriptionally active maturing oocytes potentially impedes the SUMOylation typically seen as part of the nucleolar stress response.

SUMO localized primarily in the nucleoplasm, with several depositions at sites of condensed DNA (i.e., heterochromatin) in vehicle-treated NSN-stage GV oocytes (Fig. 4C**, arrowheads**). During an examination of the effects of various types of proteotoxic stress on oocytes (Fig. 2F; **Fig. S2A**), we noticed that upon heat shock (HS; incubation at 42 °C for 2 hr), SUMO delocalized and exhibited co-localization with SC35 (a marker of nuclear speckles) -positive droplets (Fig. 4C**, NSN**). In vehicle-treated SN-stage oocytes, SUMO was highly enriched along the heterochromatin rim (Fig. 4C**, arrow**), while SC35-positive droplets disappeared. Upon HS, the SUMO signal formed SUMO droplets that tightly co-localized with SC35-positive droplets (Fig. 4C**, NSN-SN and SN**). In other experiments, we treated maturing oocytes with proteosome inhibitors for prolonged intervals. Exposure of maturing oocytes to MG132 for 39 hr (Fig. 4D; **Fig. S4A**) or to epoxomicin for 45 hr (**Fig. S4D**) resulted in the formation of large SUMO droplets that co-localized with SC35-positive droplets, consistent with the results of the HS experiments. We next tested the effect of deliberate assembly of PML-NBs on the response to proteotoxic stressors, taking into consideration the observation (data not shown) that GV oocytes were resilient to HS compared to maturing oocytes (in which the whole cells were easily distorted). However, we were unable to obtain assembly of PML-NBs in GV oocytes by injecting *hPML**VI*** mRNA (a highly effective method for driving gene expression, used here as an alternative to plasmid injection) unless the recipient oocytes were exposed to additional stimuli (**Fig. S3C,D**). Accordingly in maturing oocytes, when the exposure to a proteotoxic stressor was conducted after deliberate assembly of PML-NBs by the injection of the GFP-hPML**VI**-encoding plasmid, SUMO was sequestered not with SC35-positive droplets but with GFP-hPML**VI**-induced PML-NBs (Fig. 4E). When considered together with the data indicating that assembled PML-NBs in oocytes are not involved in the regulation of misfolded proteins upon proteotoxic stress (Fig. 2F; **Fig. S2A,B**), these results suggested that nascent PML-NBs in the nucleoplasm indirectly affect the efficiency of the SUMO response that is mediated by liquid droplet formation upon exposure to stressors.

**Fig. 4.**
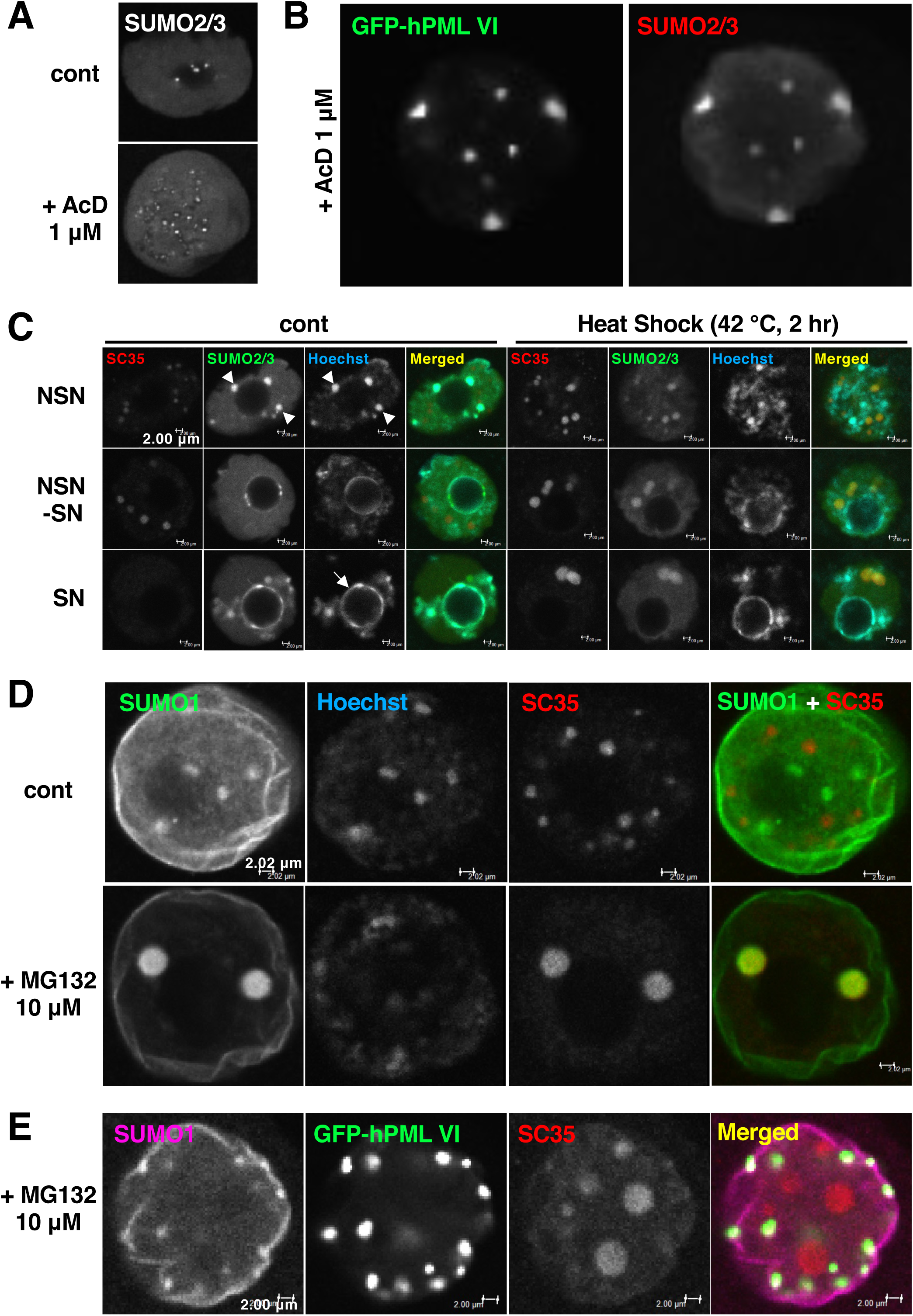
PML-NBs potentially affect the efficiency of the response of SUMO mediated by liquid droplet formation upon exposure to multiple stresses. (A) Representative images of the nuclei of maturing oocytes left untreated (cont) or treated with 1 μM actinomycin D (AcD), a transcriptional inhibitor, for 20 hr. Oocytes were stained with Alexa 488-conjugated anti-SUMO2/3 antibody. (B) Representative image of the nucleus of a maturing oocyte injected with a plasmid encoding GFP-hPML**VI** and cultured for 26 hr followed by treatment with 1 μM AcD for 21 hr. The oocyte was stained with anti-SUMO2/3 antibody (red). (C) Representative images of the nuclei of GV oocytes cultured for 17 hr and then left untreated (cont) or subjected to heat shock at 42 °C for 2 hr. NSN; non-surrounded nucleolus stage, SN; surrounded nucleolus stage. Arrowheads indicate SUMO deposition at sites of condensed DNA in NSN-oocytes. Arrow indicates the heterochromatin rim in SN-oocytes. Oocytes were stained with anti-splicing component, 35 kDa (SC35) (red) and Alexa 488-conjugated anti-SUMO2/3 (green) antibodies. Scale bars, 2.00 μm. (D) Representative images of the nuclei of maturing oocytes cultured for 26 hr and then left untreated (cont) or subjected to treatment with 10 μM MG132 for 39 hr. Prolonged exposure to proteasome inhibitors is proteotoxic to oocytes. Oocytes were stained with anti-SUMO1 (green) and anti-SC35 (red) antibodies. Scale bars, 2.02 μm. (E) An examination of whether exogenously assembled PML-NBs alter the response of SUMO to proteotoxic stress. Representative image of the nucleus of a maturing oocyte injected with a plasmid encoding GFP-hPML**VI** and cultured for 25 hr followed by treatment with 10 μM MG132 for 37 hr. Oocytes were stained with anti-SUMO1 (magenta) and anti-SC35 (red) antibodies. Scale bars, 2.00 μm.

## Discussion

We conjecture that if PML proteins were phase separated to form NBs in the nucleoplasm of oocytes, the available SUMO would be preferentially used for the maintenance of the dynamics of nascent PML-NBs. We tested this hypothesis by modifying the availability of SUMO in the nucleoplasm, both from the PML and SUMO sides. The limited number of enlarged v5-hPML**VI** (K160, 490R) -induced PML-NBs (Fig. 3B) was consistent with previous work, including studies (a) conducted in cultured cells expressing PML mutant proteins harboring the K65A mutation (K65 is another residue involved in SUMOylation, via a process that is tightly coupled to K160 SUMOylation) (Lallemand-Breitenbach et al., 2008); (b) testing PML-NBs in SUMO mutant-transfected cells that are deficient in SUMO conjugation or polymerization (Fu et al., 2005); and (c) in SUMO-conjugating enzyme (*Ubc9*) -deficient blastocysts (Nacerddine et al., 2005). A recent study employing polymer physics proposed a model for the organization of paraspeckle, another core-shell type nuclear body. That report showed that parts of the building domains of the resident (long non-coding RNAs) are mutually repulsive when positioned at the surface of paraspeckle, which accordingly determines the dynamics (size, number, and distribution) of the nuclear body; however, these repulsive domains are not required for the assembly of paraspeckle itself (Yamazaki et al., 2021). Notably, SUMOylation is dispensable for the formation of PML-NBs (Sahin et al., 2014). As shown in the wild-type v5-hPML**VI** -injected oocytes (Fig. 3C), arsenite treatment induces rapid polymerization of SUMO chains on insolubilized PML (**Fig. S3B**) at the shell, such that the structure formed spherical co-layer; this observation suggested that the structure was dominated by minimization of surface tension. The crooked-shaped shells (Fig. 3B), which are composed of mutant PML whose avidity for SUMO is limited, may indicate that the availability of SUMO in the oocyte nucleoplasm is one of the determinants of the surface property (facing the nucleoplasm) of nascent PML-NBs. Tight sequestration of SUMO molecules into the PML-negative inner core (as observed in the v5-hPML**VI** (K160, 490R) -injected oocytes) is reminiscent of the sequestration of the profluorescent biarsenical dye 4,5-bis(1,3,2-dithiarsolan-2-yl)fluorescein (FIAsH) in the cores of C212/213A-arsenic binding site mutants (Jeanne et al., 2010), and is deserving of further study. In the present work, we utilized a chemical (ML-792) to deplete the availability of SUMO in the nucleoplasm based on a mechanism of covalent SUMO-ML-792 adduct formation within the SUMO E1 enzyme (Brownell et al., 2010; He et al., 2017). We found that newly assembled GFP-hPML**VI**-induced PML-NBs frequently cluster together in maturing oocytes cultured in the presence of ML-792 (Fig. 3F). This aberrant tethering would mechanistically relate (at least in part) to the clustering of nascent v5-hPML**VI** (K160, 490R) -induced PML-NBs in the nucleoplasm upon arsenite treatment (Fig. 3D**#2**), namely arsenite-dependent boosting of SUMOylation may anyhow be attempted on mutated PML, which has little avidity for SUMO. These results suggested that a NB-free intranuclear milieu is beneficial for maturing oocytes, enhancing the availability of SUMO molecules during oocyte maturation.

Three SUMO isoforms (SUMO1, 2, and 3) are expressed ubiquitously in mammals. SUMO2 is the major isoform expressed in embryogenesis; *Sumo2*-deficient mice die at approximately embryonic day 10.5 (Wang et al., 2014). In contrast, neither SUMO1 nor SUMO3 is essential for embryogenesis. There are three distinct steps to SUMOylation; Ubc9, the sole enzyme required for the second reaction, also is indispensable for embryogenesis. Like embryos lacking SUMO2, *Ubc9*-deficient embryos die during embryogenesis, at approximately embryonic day 3.5-7.5, that is, just after implantation (Nacerddine et al., 2005). Mice harboring an oocyte-specific knockout of the *Ubc9*-encoding gene (*Ube2i-cKO*) show infertility with complete failure of oocytes to extrude the polar bodies (but no aberrancy in spindle morphology) (Rodriguez et al., 2019), supporting the notion that SUMOylation is critical for regulation of molecules involved in the MI-MII transition (Wang et al., 2010; Yuan et al., 2014; Ding et al., 2018). As has been suggested previously (Ihara et al., 2008), the *Ube2i-cKO* study indicates that the SUMOylation pathway is required not only for the resumption of meiosis but also for processes that begin before the GV stage. Since in the *Ube2i-cKO* study, Rodriguez et al. used a *Gdf-9 Cre* promotor as a driver (i.e., *Ube2i* deletion would have begun at approximately postnatal day 3), the observed dysfunction of gonadotropin-primed GV oocytes collected from 3-week-old mice implies that SUMOylation is required for a step occurring during folliculogenesis, during which oocytes undergo maturation. Although the specific relevant targets of SUMOylation currently are unknown, this previous literature suggests the significance both of SUMOylation and of SUMO availability in the nucleoplasm of maturing oocytes.

Individual maturing oocytes with maximum transcriptional activity collected from approximately postnatal day 14 gradually achieve a transcriptionally silent state, rendering these cells meiotically competent SN-type GV oocytes (De La Fuente, 2006). SN-type GV oocytes are observed from at least postnatal day 16-17 in mice and increase in proportion with mouse age (Bouniol-Baly et al., 1999; Inoue et al., 2007). Following global transcriptional repression, oocytes undergo chromosomal reconfiguration around the nucleolus to acquire the full competence to become embryos. These two events are coordinated in time, but occur independently of each other, given that *nucleoplasmin 2* (*Npm2*) -null oocytes (which fail to remodel the chromosomal configuration) still show global transcriptional repression (De La Fuente et al., 2004), and given that *mixed lineage leukemia 2* (*Mll2*) -null oocytes (which show a normal chromosomal configuration) fail to repress transcriptional activity adequately (Andreu-Vieyra et al., 2010). Oocytes mutated in these genes suffer multiple insufficiencies that affect subsequent development, suggesting that accurate regulation of these events is vital for oocyte/embryo development. In the present study, we found that SUMO, especially SUMO1, was detected as small drops in droplet (the nucleolus) of fresh maturing oocytes in which active ribosomal biogenesis involving rDNA transcription was occurring. Even though global transcriptional repression is essential for development, data from exogenously assembled PML-NBs (induced by injection of a plasmid encoding GFP-hPML**VI**) revealed that the SUMO response to AcD was impaired in such injected cells. These results suggested that, in normal oocytes devoid of PML-NBs, SUMO molecules evade sequestration by PML-NBs; this reserve of available SUMO is beneficial in case of the sudden quiescence of transcription (i.e., prematurity).

SUMO forms large droplets with SC35-positive droplets upon exposure to proteotoxic stressors. Although the droplet formation upon HS appears to accompany the arrest of enrichment of SUMO along the heterochromatin rim in SN-type GV oocytes (Fig. 4C; also note **Fig. S2C** showing heterochromatin rim in normal SN-type GV oocytes), the significance of this phenomenon remains to be determined. Whereas the enlargement of SC35-positive nuclear speckles also was observed in transcriptional repression (**Fig. S4C**), the SUMO dispersion in response to AcD (Fig. 4A), as well as the response of Ubc9 to MG132 (dissimilarly to the response to AcD) (**Fig. S4B,C**, respectively), indicated that SUMO-SC35 droplet formation upon MG132 treatment is likely independent of transcriptional repression. Together, these results suggested that SUMO, in normal oocytes devoid of PML-NBs, is involved in the acquisition of meiotic competence and participates in the emergency response to multiple stresses. Notably, the large droplets of SUMO and SC35 were no longer observed in embryos exposed to HS at approximately the early blastocyst stage, when endogenous PML-NBs emerge (**Fig. S4E**). We therefore predict that oocytes should exhibit an interval until the endogenous PML-NB emergence following the completion of oocyte development.

In conclusion, the present study demonstrated time-course-related links of PML protein to heterochromatin in the development of oocytes, during which PML does not engage in phase separation to form NBs. The PML-NB-free intranuclear milieu of oocytes reflects the significance of the reserve of SUMO available for emergent responses. The insights described here are expected to enhance our understanding of how the dynamics of membrane-less organelles contributes to cellular events and to responses to developmental cues.

## Acknowledgments

The authors thank Dr. Mounira K. Chelbi-Alix, Dr. Kenji Miyado, Dr. Azusa Inoue, and Dr. Satoshi Tsukamoto for helpful discussions. This work was supported, in part, by a Grant-in-Aid from the Japan Society for the Promotion of Science (JSPS Grant No. 16K15386, to S.H.) and research funding from the National Institute for Environmental Studies (NIES Grant No. 1620AQ026, to O.U.).

## Author Contributions

O.U. performed all experiments, with assistance from A.K-U. O.U. designed the project, analyzed data, prepared figures, and wrote the manuscript. S.H. supervised the project and wrote the manuscript.

## Declaration of Interests

The authors declare they have no actual or potential competing financial interests.

## Figure legends

**Fig. S1, related to Fig. 1. Subcellular localization of endogenous PML in metaphase-stage oocytes.** (A) Representative electron micrograph image of peri-chromosomal localization of endogenous PML visualized with primary anti-mouse PML (mPML) antibody and gold particle-conjugated (Gold) secondary antibody. Germinal vesicle (GV) oocytes were cultured *in vitro* for 20 hr. A magnified electron micrograph is shown on the right. (B-D) Representative fluorescent images of oocytes collected from the ovaries of adult mice. The chromosome arm of the metaphase II (MII) -stage oocyte (B, z-stack), the kinetochore of the GV breakdown (GVBD)- metaphase I (MI) -stage oocyte (C, z-stack), and the early endosome of the MII-stage oocyte (D) were labeled with anti-survivin antibody, calcium-responsive transactivator (CREST) antiserum, and anti-early endosome antigen 1 (EEA1) antibody, respectively. BF, bright-field image of the oocyte. (E) Representative image of an embryo at approximately the early blastocyst stage. (Upper) Subcellular localization of endogenous PML (red) and SUMO2/3 (green). Scale bars, 975 nm. (Lower) Co-localization of SUMO2/3 (green) with DAXX (red), a representative client of PML-NBs. Images were reconstructed as z-stack images. Scale bars, 20.2 µm. (F) Representative image of endogenous PML-NBs (red) in control bone marrow-derived cells.

**Fig. S2, related to Fig. 2. An examination of whether exogenously assembled and endogenously appearing PML-NBs act as overflow compartments for misfolded proteins.** (A) Representative image of the nucleus of a maturing oocyte injected with a plasmid encoding GFP-hPML**VI** and cultured for the indicated time. Before the end of culturing, newly synthesized aberrant polypeptides were labeled with 20 μM OP-puro (red) combined with 100 μM MG132 for the indicated time. BF, bright-field images of the oocytes. Scale bars, 2.00 μm. (B) (Left) Representative image of an embryo cultured for 76 hr before labeling of newly synthesized aberrant polypeptides with 20 μM OP-puro (red) combined with 10 μM MG132 for 4 hr (80 hr post-insemination, 80 h.p.i.). Endogenously appearing PML-NBs in the nucleus of each blastomere at the morula to early blastocyst stage were visualized by immunofluorescent staining with anti-mouse PML (mPML, green) antibody. BF, bright-field image of the embryo. Scale bars, 19.9 μm. (Right) Enlarged images of the nuclei of representative blastomeres. Scale bars, 2.00 µm. (C) Representative image of the nucleus of a germinal vesicle (GV) oocyte treated with dimethyl sulfoxide (DMSO, vehicle control) for 24 hr. Endogenous PML was visualized by staining with anti-mouse PML (mPML, red) antibody. The oocyte was further stained with Alexa 488-conjugated anti-SUMO2/3 antibody.

**Fig. S3, related to Fig. 3. Preparations for modifying the SUMO availability in oocytes.** (A) Schematic domain structures of wild-type human PML**VI** and a mutant version of the same protein that has limited avidity for SUMO; these proteins were encoded by the plasmids v5-hPML**VI** and v5-hPML**VI** (K160, 490R), respectively. RING: Really Interesting New Gene domain, with conserved cysteine. B1 and B2: zinc-binding boxes. CC: coiled-coil domain. Two SUMOylation sites are indicated by aqua blue circles labeled with an S. These sites were mutated from lysine to arginine in the SUMOylation-deficient mutant protein encoded by the v5-hPML**VI** (K160, 490R) construct. (B) Expression of constructs and biochemical response of their products to arsenite. Immunoblot analysis of CHO-K1 cells transiently expressing wild-type human PML**VI** or the SUMOylation-deficient mutant protein. The fractions soluble or insoluble to radioimmunoprecipitation (RIPA) lysis buffer were immunoblotted (IB) with the indicated antibodies. The cells were exposed to 3 μM arsenite (As) for 3 hr. Green asterisk indicates the biochemical response of the v5-hPML**VI** (K160, 490R)-encoded mutant protein to arsenite (i.e., corresponding to a shift into the detergent-resistant nuclear matrix (Lallemand-Breitenbach et al., 2008)), which was comparable to that seen with the unmutated protein encoded by the v5-hPML**VI** construct. (C and D) Representative images of the nuclei of GV oocytes injected with the mRNA transcribed *in vitro* from plasmid v5-hPML**VI** (encoding wild-type human PML**VI**) (C) or v5-hPML**VI** (K160, 490R) (encoding human PML**VI** with limited avidity for SUMO) (D) and cultured for 46 hr or 48 hr, respectively, and then left untreated (cont) or subjected to treatment with 3 μM arsenite for 2 hr. Oocytes were stained with anti-human PML (red) and Alexa 488-conjugated anti-SUMO2/3 (green) antibodies. NSN; non-surrounded nucleolus stage, SN; surrounded nucleolus stage. Scale bars, 2.00 μm.

**Fig. S4, related to Fig. 4. Characterization of SUMO-SC35 droplets.** (A and B) Representative images of the nuclei of maturing oocytes treated with dimethyl sulfoxide (DMSO, vehicle control) or 20 μM ML-792 or 10 μM MG132 for the indicated time. (A) Oocytes were stained with anti-SC35 (red) and Alexa 488-conjugated anti-SUMO2/3 (green) antibodies. (B) Oocytes were stained with anti-Ubc9 antibody. (C) Comparison of the response of SC35 droplets with and without transcriptional inhibition. Representative images of the nuclei of maturing oocytes treated with DMSO or 1 μM actinomycin D (AcD) for 16 hr. Oocytes were stained with anti-SC35 (red) and anti-Ubc9 (green) antibodies. Scale bars, 2.00 μm. (D) Representative z-stack image of a maturing oocyte cultured in the presence of 1 μM epoxomicin (Epox), another proteosome inhibitor, for 45 hr. Oocytes were stained with anti-SUMO1 antibody. An enlarged image of the nucleus is shown at the bottom. BF, bright-field image of the oocyte. Scale bars, 19.9 μm. (E) Representative images of post-morula-stage embryos. Embryos at 80 hr post-insemination (80 h.p.i.) subsequently were left untreated (cont) or exposed to heat shock at 42 °C for 2 hr. Embryos were stained with anti-SC35 (red) and Alexa 488-conjugated anti-SUMO2/3 (green) antibodies. BF, bright-field images of the embryos. Scale bars, 19.9 μm.

## Notes

### Competing Interest Statement

The authors have declared no competing interest.

## References

1. Andreu-Vieyra, C.V., Chen, R., Agno, J.E., Glaser, S., Anastassiadis, K., Stewart, A.F., and Matzuk, M.M. (2010). MLL2 is required in oocytes for bulk histone 3 lysine 4 trimethylation and transcriptional silencing. PLoS biology 8.

2. Banani, S.F., Lee, H.O., Hyman, A.A., and Rosen, M.K. (2017). Biomolecular condensates: organizers of cellular biochemistry. Nat Rev Mol Cell Biol 18, 285–298

3. Banani, S.F., Rice, A.M., Peeples, W.B., Lin, Y., Jain, S., Parker, R., and Rosen, M.K. (2016). Compositional Control of Phase-Separated Cellular Bodies. Cell 166, 651–663.

4. Bernardi, R., and Pandolfi, P.P. (2003). Role of PML and the PML-nuclear body in the control of programmed cell death. Oncogene 22, 9048–9057.

5. Bernardi, R., Scaglioni, P.P., Bergmann, S., Horn, H.F., Vousden, K.H., and Pandolfi, P.P. (2004). PML regulates p53 stability by sequestering Mdm2 to the nucleolus. Nature cell biology 6, 665–672.

6. Bersaglieri, C., and Santoro, R. (2019). Genome Organization in and around the Nucleolus. Cells 8.

7. Bøe, S.O., Haave, M., Jul-Larsen, Å., Grudic, A., Bjerkvig, R., and Lønning, P.E. (2006). Promyelocytic leukemia nuclear bodies are predetermined processing sites for damaged DNA. Journal of cell science 119, 3284–3295.

8. Bouniol-Baly, C., Hamraoui, L., Guibert, J., Beaujean, N., Szöllösi, M.S., and Debey, P. (1999). Differential transcriptional activity associated with chromatin configuration in fully grown mouse germinal vesicle oocytes. Biology of Reproduction 60, 580–587.

9. Brownell, J.E., Sintchak, M.D., Gavin, J.M., Liao, H., Bruzzese, F.J., Bump, N.J., Soucy, T.A., Milhollen, M.A., Yang, X., Burkhardt, A.L., et al. (2010). Substrate-assisted inhibition of ubiquitin-like protein-activating enzymes: the NEDD8 E1 inhibitor MLN4924 forms a NEDD8-AMP mimetic in situ. Mol Cell 37, 102–111.

10. Cappadocia, L., Mascle, X.H., Bourdeau, V., Tremblay-Belzile, S., Chaker-Margot, M., Lussier-Price, M., Wada, J., Sakaguchi, K., Aubry, M., Ferbeyre, G., et al. (2015). Structural and functional characterization of the phosphorylation-dependent interaction between PML and SUMO1. Structure 23, 126–138.

11. Chelbi-Alix, M., Pelicano, L., Quignon, F., Koken, M., Venturini, L., Stadler, M., Pavlovic, J., and Degos, L. (1995). Induction of the PML protein by interferons in normal and APL cells. Leukemia 9, 2027–2033.

12. Cho, S., Park, J.S., and Kang, Y.-K. (2011). Dual Functions of Histone-Lysine N-Methyltransferase Setdb1 Protein at Promyelocytic Leukemia-Nuclear Body (PML-NB) MAINTAINING PML-NB STRUCTURE AND REGULATING THE EXPRESSION OF ITS ASSOCIATED GENES. Journal of Biological Chemistry 286, 41115–41124.

13. De La Fuente, R. (2006). Chromatin modifications in the germinal vesicle (GV) of mammalian oocytes. Developmental biology 292, 1–12.

14. De La Fuente, R., Viveiros, M.M., Burns, K.H., Adashi, E.Y., Matzuk, M.M., and Eppig, J.J. (2004). Major chromatin remodeling in the germinal vesicle (GV) of mammalian oocytes is dispensable for global transcriptional silencing but required for centromeric heterochromatin function. Developmental biology 275, 447–458.

15. Dellaire, G., Eskiw, C.H., Dehghani, H., Ching, R.W., and Bazett-Jones, D.P. (2006). Mitotic accumulations of PML protein contribute to the re-establishment of PML nuclear bodies in G1. Journal of cell science 119, 1034–1042.

16. Ding, Y., Kaido, M., Llano, E., Pendas, A.M., and Kitajima, T.S. (2018). The Post-anaphase SUMO Pathway Ensures the Maintenance of Centromeric Cohesion through Meiosis I-II Transition in Mammalian Oocytes. Curr Biol 28, 1661–1669 e1664.

17. Draskovic, I., Arnoult, N., Steiner, V., Bacchetti, S., Lomonte, P., and Londoño-Vallejo, A. (2009). Probing PML body function in ALT cells reveals spatiotemporal requirements for telomere recombination. Proceedings of the National Academy of Sciences 106, 15726–15731.

18. Ebrahimian, M., Mojtahedzadeh, M., Bazett-Jones, D., and Dehghani, H. (2010). Transcript isoforms of promyelocytic leukemia in mouse male and female gametes. Cells Tissues Organs 192, 374–381.

19. Everett, R.D., and Chelbi-Alix, M.K. (2007). PML and PML nuclear bodies: implications in antiviral defence. Biochimie 89, 819–830.

20. Ferbeyre, G., de Stanchina, E., Querido, E., Baptiste, N., Prives, C., and Lowe, S.W. (2000). PML is induced by oncogenic ras and promotes premature senescence. Genes & development 14, 2015–2027.

21. Feric, M., Vaidya, N., Harmon, T.S., Mitrea, D.M., Zhu, L., Richardson, T.M., Kriwacki, R.W., Pappu, R.V., and Brangwynne, C.P. (2016). Coexisting Liquid Phases Underlie Nucleolar Subcompartments. Cell 165, 1686–1697.

22. Flemr, M., Ma, J., Schultz, R.M., and Svoboda, P. (2010). P-body loss is concomitant with formation of a messenger RNA storage domain in mouse oocytes. Biol Reprod 82, 1008–1017.

23. Flynn, R.L., Cox, K.E., Jeitany, M., Wakimoto, H., Bryll, A.R., Ganem, N.J., Bersani, F., Pineda, J.R., Suvà, M.L., and Benes, C.H. (2015). Alternative lengthening of telomeres renders cancer cells hypersensitive to ATR inhibitors. Science 347, 273–277.

24. Fu, C., Ahmed, K., Ding, H., Ding, X., Lan, J., Yang, Z., Miao, Y., Zhu, Y., Shi, Y., and Zhu, J. (2005). Stabilization of PML nuclear localization by conjugation and oligomerization of SUMO-3. Oncogene 24, 5401–5413.

25. Fulka, H., and Aoki, F. (2016). Nucleolus Precursor Bodies and Ribosome Biogenesis in Early Mammalian Embryos: Old Theories and New Discoveries. Biol Reprod 94, 143

26. Fulka, H., Rychtarova, J., and Loi, P. (2020). The nucleolus-like and precursor bodies of mammalian oocytes and embryos and their possible role in post-fertilization centromere remodelling. Biochem Soc Trans 48, 581–593.

27. Goddard, A., Yuan, J., Fairbairn, L., Dexter, M., Borrow, J., Kozak, C., and Solomon, E. (1995). Cloning of the murine homolog of the leukemia-associated PML gene. Mammalian Genome 6, 732–737.

28. He, X., Riceberg, J., Soucy, T., Koenig, E., Minissale, J., Gallery, M., Bernard, H., Yang, X., Liao, H., Rabino, C., et al. (2017). Probing the roles of SUMOylation in cancer cell biology by using a selective SAE inhibitor. Nat Chem Biol 13, 1164–1171.

29. Hirano, S., Tadano, M., Kobayashi, Y., Udagawa, O., and Kato, A. (2015). Solubility shift and SUMOylaltion of promyelocytic leukemia (PML) protein in response to arsenic(III) and fate of the SUMOylated PML. Toxicol Appl Pharmacol 287, 191–201.

30. Hirano, S., Udagawa, O., Kobayashi, Y., and Kato, A. (2018). Solubility changes of promyelocytic leukemia (PML) and SUMO monomers and dynamics of PML nuclear body proteins in arsenite-treated cells. Toxicology and applied pharmacology 360, 150–159.

31. Ihara, M., Stein, P., and Schultz, R.M. (2008). UBE2I (UBC9), a SUMO-conjugating enzyme, localizes to nuclear speckles and stimulates transcription in mouse oocytes. Biol Reprod 79, 906–913.

32. Inoue, A., Akiyama, T., Nagata, M., and Aoki, F. (2007). The perivitelline space-forming capacity of mouse oocytes is associated with meiotic competence. Journal of Reproduction and Development 53, 1043–1052.

33. Jeanne, M., Lallemand-Breitenbach, V., Ferhi, O., Koken, M., Le Bras, M., Duffort, S., Peres, L., Berthier, C., Soilihi, H., Raught, B., et al. (2010). PML/RARA oxidation and arsenic binding initiate the antileukemia response of As2O3. Cancer Cell 18, 88–98.

34. Lallemand-Breitenbach, V., and de The, H. (2018). PML nuclear bodies: from architecture to function. Curr Opin Cell Biol 52, 154–161.

35. Lallemand-Breitenbach, V., Jeanne, M., Benhenda, S., Nasr, R., Lei, M., Peres, L., Zhou, J., Zhu, J., Raught, B., and de Thé, H. (2008). Arsenic degrades PML or PML–RARα through a SUMO-triggered RNF4/ubiquitin-mediated pathway. Nature cell biology 10, 547–555.

36. Lallemand-Breitenbach, V., Zhu, J., Puvion, F., Koken, M., Honoré, N., Doubeikovsky, A., Duprez, E., Pandolfi, P.P., Puvion, E., and Freemont, P. (2001). Role of promyelocytic leukemia (PML) sumolation in nuclear body formation, 11S proteasome recruitment, and As2O3-induced PML or PML/retinoic acid receptor α degradation. The Journal of experimental medicine 193, 1361–1372.

37. Lamond, A.I., and Earnshaw, W.C. (1998). Structure and function in the nucleus. Science 280, 547–553.

38. Li, Y., Ma, X., Chen, Z., Wu, H., Wang, P., Wu, W., Cheng, N., Zeng, L., Zhang, H., Cai, X., et al. (2019). B1 oligomerization regulates PML nuclear body biogenesis and leukemogenesis. Nat Commun 10, 3789.

39. Louria-Hayon, I., Grossman, T., Sionov, R.V., Alsheich, O., Pandolfi, P.P., and Haupt, Y. (2003). The promyelocytic leukemia protein protects p53 from Mdm2-mediated inhibition and degradation. Journal of Biological Chemistry 278, 33134–33141.

40. Lunardi, A., Gaboli, M., Giorgio, M., Rivi, R., Bygrave, A., Antoniou, M., Drabek, D., Dzierzak, E., Fagioli, M., and Salmena, L. (2011). A role for PML in innate immunity. Genes & cancer 2, 10–19.

41. Mediani, L., Guillen-Boixet, J., Vinet, J., Franzmann, T.M., Bigi, I., Mateju, D., Carra, A.D., Morelli, F.F., Tiago, T., Poser, I., et al. (2019). Defective ribosomal products challenge nuclear function by impairing nuclear condensate dynamics and immobilizing ubiquitin. EMBO J 38, e101341.

42. Mu□ller, S., Miller Jr, W.H., and Dejean, A. (1998). Trivalent Antimonials Induce Degradation of the PML-RARα Oncoprotein and Reorganization of the Promyelocytic Leukemia Nuclear Bodies in Acute Promyelocytic Leukemia NB4 Cells. Blood, The Journal of the American Society of Hematology 92, 4308–4316.

43. Nacerddine, K., Lehembre, F., Bhaumik, M., Artus, J., Cohen-Tannoudji, M., Babinet, C., Pandolfi, P.P., and Dejean, A. (2005). The SUMO pathway is essential for nuclear integrity and chromosome segregation in mice. Developmental cell 9, 769–779.

44. Nisole, S., Maroui, M.A., Mascle, X., Aubry, M., and Chelbi-Alix, M.K. (2013). Differential roles of PML isoforms. Frontiers in oncology 3, 125.

45. Niwa-Kawakita, M., Ferhi, O., Soilihi, H., Le Bras, M., Lallemand-Breitenbach, V., and de Thé, H. (2017). PML is a ROS sensor activating p53 upon oxidative stress. Journal of Experimental Medicine 214, 3197–3206.

46. Pearson, M., Carbone, R., Sebastiani, C., Cioce, M., Fagioli, M., Saito, S.i., Higashimoto, Y., Appella, E., Minucci, S., and Pandolfi, P.P. (2000). PML regulates p53 acetylation and premature senescence induced by oncogenic Ras. Nature 406, 207–210.

47. Puvion-Dutilleul, F., Chelbi-Alix, M.K., Koken, M., Quignon, F., Puvion, E., and De Thé, H. (1995). Adenovirus infection induces rearrangements in the intranuclear distribution of the nuclear body-associated PML protein. Experimental cell research 218, 9–16.

48. Racki, W.J., and Richter, J.D. (2006). CPEB controls oocyte growth and follicle development in the mouse. Development 133, 4527–4537.

49. Rodriguez, A., Briley, S.M., Patton, B.K., Tripurani, S.K., Rajapakshe, K., Coarfa, C., Rajkovic, A., Andrieux, A., Dejean, A., and Pangas, S.A. (2019). Loss of the E2 SUMO-conjugating enzyme Ube2i in oocytes during ovarian folliculogenesis causes infertility in mice. Development 146.

50. Sahin, U., Ferhi, O., Jeanne, M., Benhenda, S., Berthier, C., Jollivet, F., Niwa-Kawakita, M., Faklaris, O., Setterblad, N., de The, H., et al. (2014). Oxidative stress-induced assembly of PML nuclear bodies controls sumoylation of partner proteins. J Cell Biol 204, 931–945.

51. Sundqvist, A., Liu, G., Mirsaliotis, A., and Xirodimas, D.P. (2009). Regulation of nucleolar signalling to p53 through NEDDylation of L11. EMBO reports 10, 1132–1139.

52. Uozumi, N., Matsumoto, H., and Saitoh, H. (2016). Detection of O-propargyl-puromycin with SUMO and ubiquitin by click chemistry at PML-nuclear bodies during abortive proteasome activities. Biochem Biophys Res Commun 474, 247–251.

53. Wang, L., Wansleeben, C., Zhao, S., Miao, P., Paschen, W., and Yang, W. (2014). SUMO2 is essential while SUMO3 is dispensable for mouse embryonic development. EMBO reports 15, 878–885.

54. Wang, P., Benhenda, S., Wu, H., Lallemand-Breitenbach, V., Zhen, T., Jollivet, F., Peres, L., Li, Y., Chen, S.J., Chen, Z., et al. (2018). RING tetramerization is required for nuclear body biogenesis and PML sumoylation. Nat Commun 9, 1277.

55. Wang, Z.B., Ou, X.H., Tong, J.S., Li, S., Wei, L., Ouyang, Y.C., Hou, Y., Schatten, H., and Sun, Q.Y. (2010). The SUMO pathway functions in mouse oocyte maturation. Cell Cycle 9, 2640–2646.

56. Wang, Z.G., Delva, L., Gaboli, M., Rivi, R., Giorgio, M., Cordon-Cardo, C., Grosveld, F., and Pandolfi, P.P. (1998). Role of PML in cell growth and the retinoic acid pathway. Science 279, 1547–1551.

57. Yamazaki, T., Yamamoto, T., Yoshino, H., Souquere, S., Nakagawa, S., Pierron, G., and Hirose, T. (2021). Paraspeckles are constructed as block copolymer micelles. EMBO J 40, e107270.

58. Yuan, Y.F., Zhai, R., Liu, X.M., Khan, H.A., Zhen, Y.H., and Huo, L.J. (2014). SUMO-1 plays crucial roles for spindle organization, chromosome congression, and chromosome segregation during mouse oocyte meiotic maturation. Mol Reprod Dev 81, 712–724.

