## Supplementary figures and images for "Promyelocytic leukemia nuclear body (PML-NB) -free intranuclear milieu facilitates development of oocytes in mice"

### Supplementary Figures

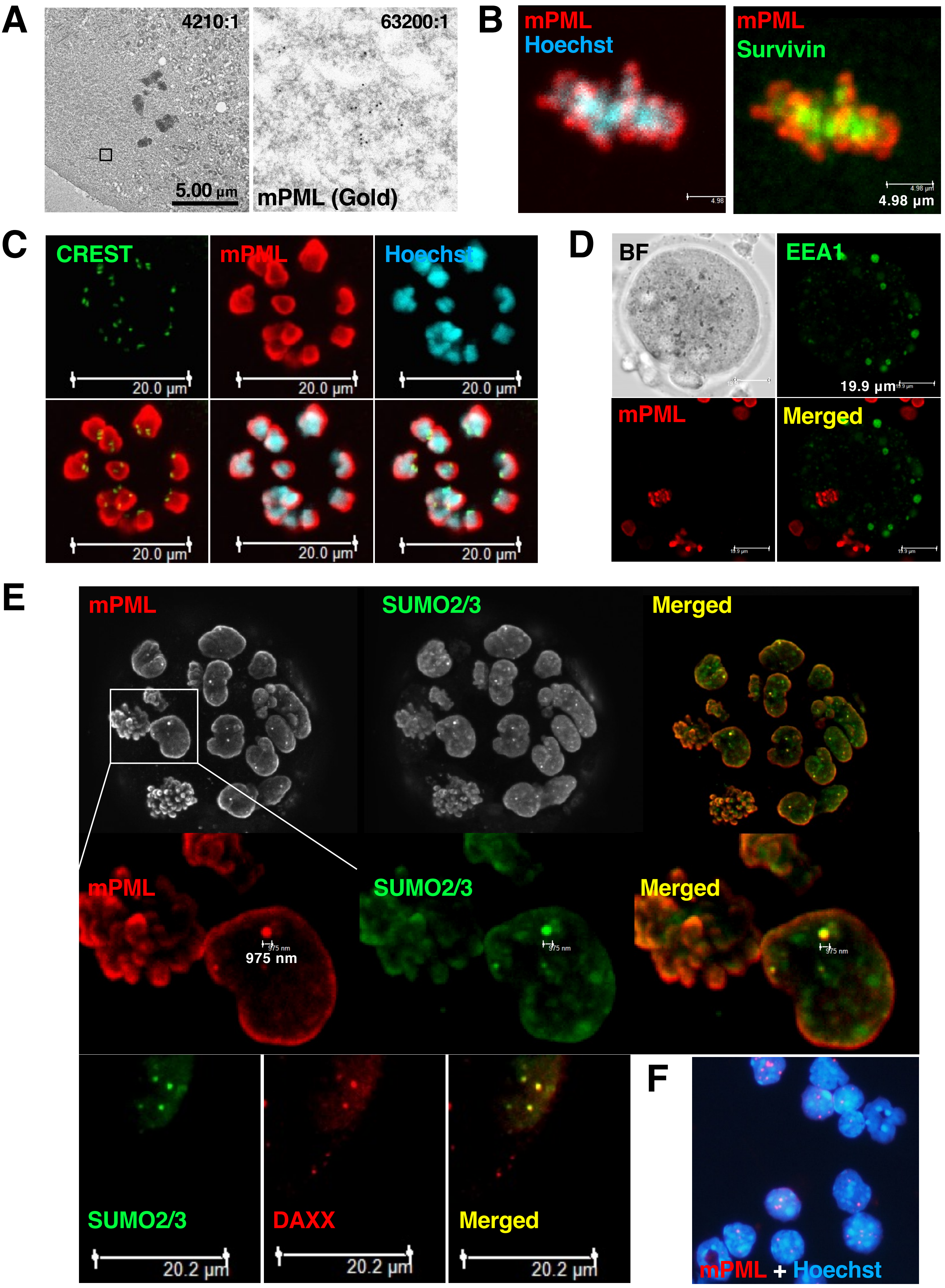

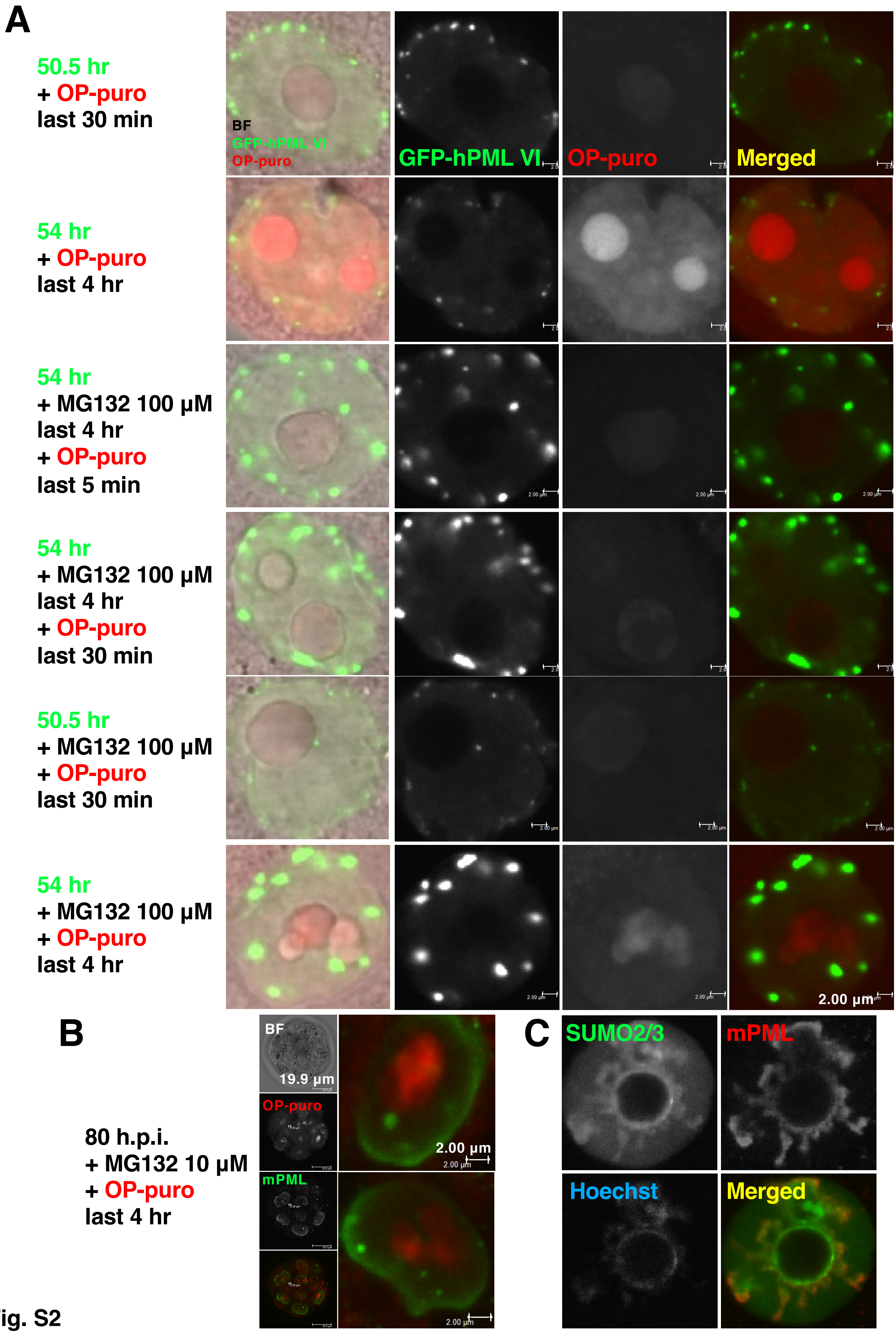

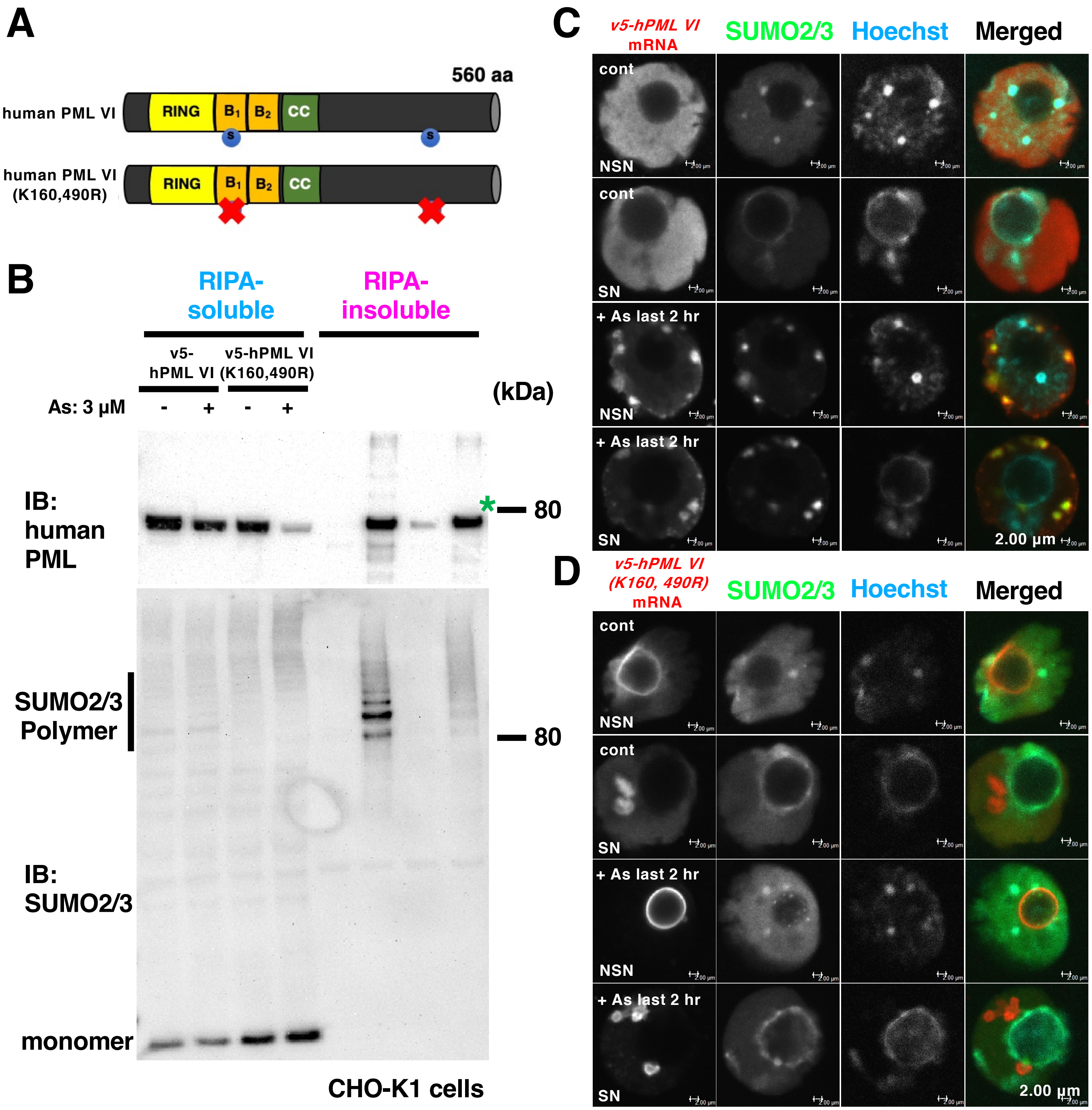

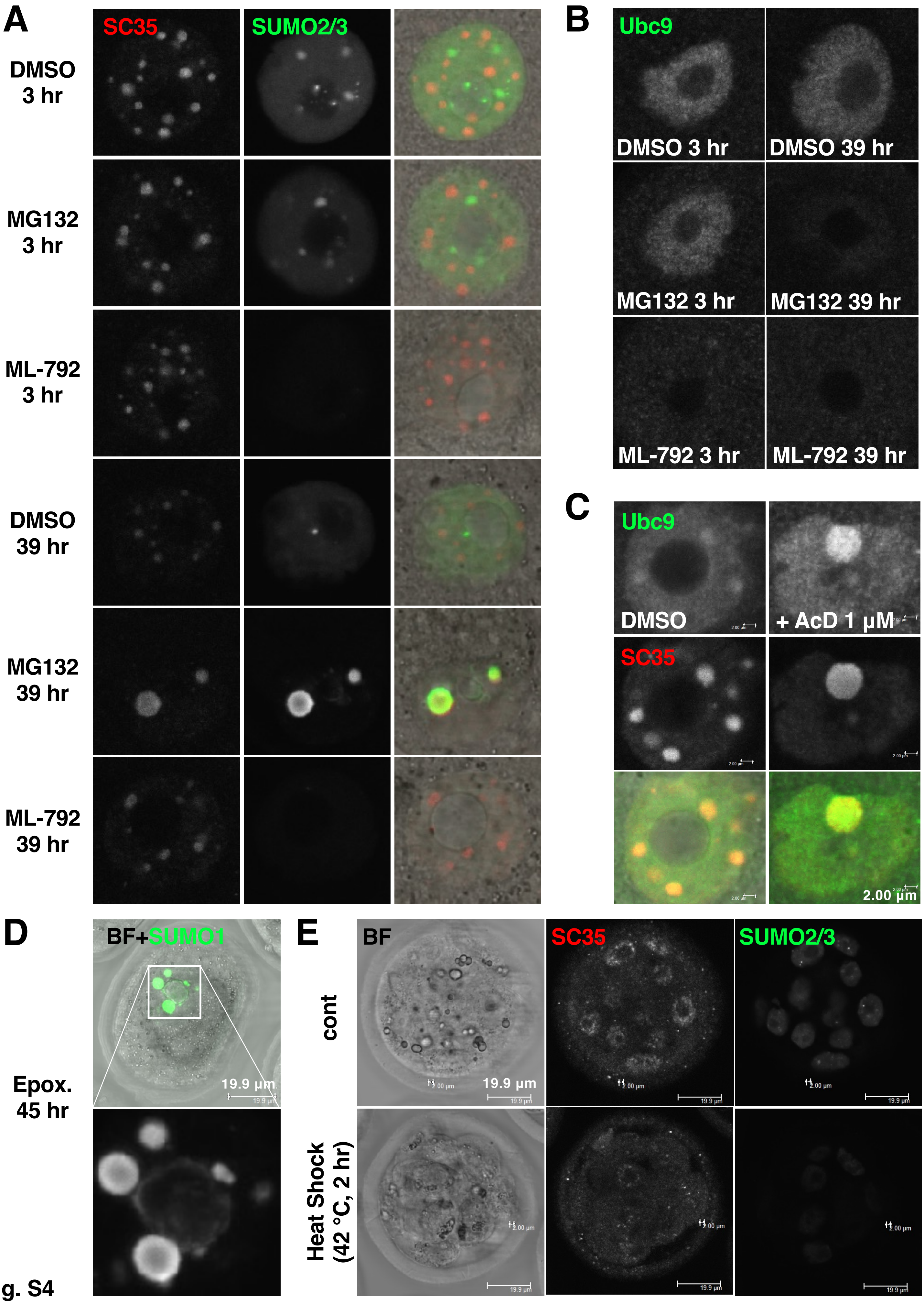
